# Antimicrobial resistance of *Enterococcus cecorum*: ECOFF determination

**DOI:** 10.1101/2022.10.19.512977

**Authors:** Jeanne Laurentie, Gwenaelle Mourand, Pauline Grippon, Sylviane Furlan, Claire Chauvin, Eric Jouy, Pascale Serror, Isabelle Kempf

**Affiliations:** ANSES, Laboratoire de Ploufragan-Plouzané-Niort, Ploufragan, France; Université Paris-Saclay, INRAE, AgroParisTech, Micalis Institute, 78350 Jouy en Josas, France

**Keywords:** *Enterococcus cecorum*, epidemiological cut-offs, disc diffusion, broth microdilution, resistance genes, MICs

## Abstract

*Enterococcus cecorum*, a commensal Gram-positive bacterium of the chicken gut, has emerged as a worldwide cause of lameness in poultry, particularly in fast-growing broilers. It is responsible for osteomyelitis, spondylitis and femoral head necrosis, causing animal suffering, mortality and antimicrobial use. Research on the antimicrobial resistance of *E. cecorum* clinical isolates in France is scarce, and epidemiological cut-off (ECOFF) values unknown. To determine tentative ECOFF (CO_WT_) values for *E. cecorum* and to investigate the antimicrobial resistance patterns of isolates from mainly French broilers, we tested the susceptibility of a collection of commensal and clinical isolates (n=208) to 29 antimicrobials by the disc diffusion (DD) method. We also determined the minimum inhibitory concentrations (MICs) of 23 antimicrobials by the broth micro-dilution method. To detect chromosomal mutations conferring antimicrobial resistance, we investigated the genomes of 118 *E. cecorum* isolates mainly obtained from infectious sites and previously described in the literature. We determined the CO_WT_ values for more than 20 antimicrobials and identified two chromosomal mutations explaining fluoroquinolone resistance. The DD method appears better suited for detecting *E. cecorum* antimicrobial resistance. Although tetracycline and erythromycin resistances were persistent in clinical and non-clinical isolates, we found little or no resistance to medically important antimicrobials.

## INTRODUCTION

*Enterococcus cecorum*, first described as *Streptococcus cecorum* in 1983 (1), is a dominant commensal Gram-positive bacterium found in the intestinal microbiota of healthy adult chickens (1). Since 2002, it has been emerging in poultry production as a pathogen responsible for lameness, osteomyelitis and spondylitis with increased mortality rates in broiler productions and to a lesser extent in ducks and other avian species (2). The development of *E. cecorum* infections is multifactorial, and depends on predisposing factors related to host genetics, rapid growth, feed composition, husbandry procedures and animal density, all in combination with the bacterium’s pathogenic potential (3-5).

There are no vaccines against *E. cecorum*, and protective measures can only include biosecurity and good poultry management. If an *E. cecorum* infection is diagnosed, antimicrobials may be used as soon as possible to prevent further progression (4). Indeed, antimicrobial therapy is ineffective for paralysed birds. The choice of the antimicrobial compound to be used should be based on the susceptibility of the isolated strain. As defined by EUCAST (https://mic.eucast.org/), epidemiological cut-off (ECOFF) values distinguish microorganisms without (wild type) and with phenotypically detectable acquired resistance mechanisms (non-wild type) to the agent in question, and a micro-organism is defined as wild type (WT) for a species by the absence of acquired and mutational resistance mechanisms to the drug in question. On the other hand, clinical breakpoints are determined on the basis of dosages, pharmacokinetics, resistance mechanisms, MIC distributions, inhibition zone diameters and more recently, pharmacodynamics and ECOFFs. A micro-organism is thus defined as clinically susceptible by a level of antimicrobial activity associated with a high likelihood of therapeutic success. Despite the few MIC criteria for *E. cecorum* proposed by Borst et al. (6) for several antimicrobials, there are no ECOFFs nor clinical breakpoints specifically for *E. cecorum*. Few studies on the antimicrobial resistance of *E. cecorum* are available and interpretation is usually based on enterococci ECOFFs or breakpoints (6-9).

Resistance to tetracycline and erythromycin is the most commonly reported, but resistance to aminoglycosides and β-lactams is sometimes detected (4, 9). To date, few acquired antimicrobial resistance genes (9) and no resistance-conferring mutations have been associated with antimicrobial resistance phenotypes in *E. cecorum*.

The aims of this study were to analyse by disc diffusion (DD) and broth micro-dilution (BMD) methods a large collection of commensal and clinical *E. cecorum* isolates to: (i) establish provisional ECOFF (CO_WT_) values for *E. cecorum* inhibition zone diameters and minimum inhibitory concentrations (MICs); (ii) to evaluate the correlation between inhibition zone diameters and MIC values; and (iii) to analyse the relationship between observed phenotypes and the genotypes of previously studied *E. cecorum* clinical isolates (J. Laurentie, V. Loux, C. Hennequet-Antier, E. Chambellon, J. Deschamps, A. Trotereau, S. Furlan, C. Darrigo, F. Kempf, J. Lao, M. Milhes, C. Roques, B. Quinquis, C. Vandecasteele, R. Boyer, O. Bouchez, F. Repoila, J. Le Guennec, H. Chiapello, R. Briandet, E. Helloin, C. Schouler, I. Kempf, and P. Serror, submitted for publication).

## MATERIAL AND METHODS

### Bacterial Isolates

The present study investigated 208 isolates of *E. cecorum*. Of these, 118 were previously described in the literature isolates (J. Laurentie, V. Loux, C. Hennequet-Antier, E. Chambellon, J. Deschamps, A. Trotereau, S. Furlan, C. Darrigo, F. Kempf, J. Lao, M. Milhes, C. Roques, B. Quinquis, C. Vandecasteele, R. Boyer, O. Bouchez, F. Repoila, J. Le Guennec, H. Chiapello, R. Briandet, E. Helloin, C. Schouler, I. Kempf, and P. Serror, submitted for publication), five were co-isolated with *E. coli* from an infected bird, and 85 were isolated between 2019 and 2021 in France from the caecal contents or pooled faecal samples of healthy birds collected from hatcheries, farms and slaughterhouses (**Table S1**). A selective medium specific to *E. cecorum* was used to isolate commensal strains (S. Furlan & P. Serror, personal communication).

### Disc diffusion method

The DD method was applied to 29 antimicrobials or associations of antimicrobials of importance in either veterinary or human medicine (amoxicillin 25 μg, ampicillin 2 μg, cefotaxime 5 μg, ceftaroline 5 μg, imipenem 10 μg, norfloxacin 10 μg, ciprofloxacin 5 μg, levofloxacin 5 μg, vancomycin 5 μg, teicoplanin 30 μg, gentamicin 500 μg, streptomycin 300 μg, spectinomycin 100 μg, lincomycin-spectinomycin 2 μg/100 μg, lincomycin 15 μg, erythromycin 15 μg, spiramycin 100 μg, tylosin 30 μg, quinupristin-dalfopristin 15 μg, tetracycline 30 μg, doxycycline 30 μg, tigecycline 15 μg, tiamulin 30 μg, fosfomycin 200 μg, nitrofurantoin 100 μg, linezolid 10 μg, chloramphenicol 30 μg, bacitracin 130 μg and rifampicin 5 μg) on Mueller-Hinton agar plates supplemented with 5% mechanically defibrinated horse blood and 20 mg/L β-NAD (MH-F agar). All the procedures were in keeping with the EUCAST DD method (10). Plates were inoculated by swabbing in three directions with a 0.5 McFarland bacterial suspension (5 × 10^7^ colony forming units (CFU)/mL), prepared in saline solution (0.85% NaCl). Inhibition zone diameters were measured after 18 h ± 2 h of incubation (and 24 h more for glycopeptides) at 35°C ± 2°C in 5% CO_2_, as recommended for S*treptococcus pneumoniae*. As there is no recommended reference strain for *E. cecorum, S. pneumoniae* (CIP 104340) was used as a proxy.

### Broth micro-dilution method

The BMD method (11) was used to determine the MICs of 23 antimicrobial agents for *E. cecorum* isolates placed in 96-well microtitre plates. Firstly, 50 μL of sterile water was placed in each well, then 50 μL of the antimicrobial solution was added to each well in the first column of the microtitre plate. Serial two-fold dilutions of the antimicrobial solutions were performed by transferring 50 μL from the first column to the second one, and subsequently up to the lowest tested dilution as described in **Figure 1**. The microtitre plates were stored at -70°C before use.

**Figure 1:**
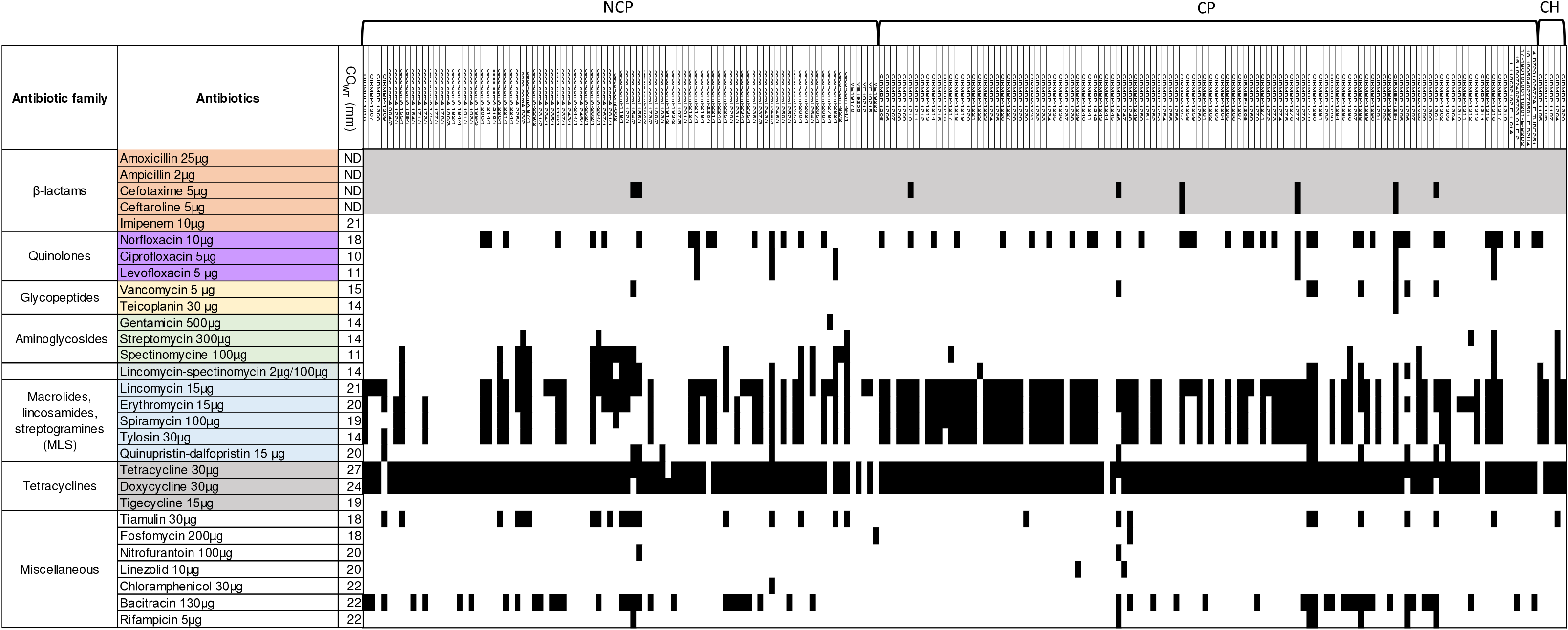
MIC results and CO_WT_ determination. White fields represent the range of dilutions tested. MIC values equal to or lower than the lowest concentration tested are presented as the lowest concentration. MIC values greater than the highest concentration tested are presented as one dilution step above the test range. A black vertical line indicates that the CO_WT_ value was calculated with the ECOFFinder tool. A dotted line indicates that the CO_WT_ value was determined by visual inspection (not enough WT isolates to calculate it). No CO_WT_ could be determined for ampicillin, amoxicillin and spectinomycin. ^a^ Quinu-Dalfo: Quinupristin-Dalfopristin

The day before the MIC determination assay, *E. cecorum* isolates were inoculated onto Columbia agar supplemented with 5% of sheep blood from Bio-Rad and incubated at 35°C for 24 hours in 5% CO_2_. Next, 0.5 McFarland bacterial suspensions prepared in saline solution were diluted 1:100 in cation-adjusted Mueller-Hinton fastidious (MH-F) broth (Thermo Scientific™) in order to reach the final concentration of 5 × 10^5^ CFU/mL. Finally, 50 μL of the suspension was added to each well. One well per plate was used as a positive control (wells with only the bacterial suspension) and one as a negative control (wells with only sterile cation-adjusted MH-F broth used to prepare the inoculum). The MICs were read after 18 ± 2 h of incubation at 35 ± 2°C and 24 h more for streptomycin, as recommended for enterococci. As there is no reference for *E. cecorum, S. pneumoniae* (CIP 104340) was used as a proxy.

### Cut-off value determination and statistical tests

For the DD method, the normalised resistance interpretation (NRI) method was used with permission from the patent holder, Bioscand AB, Täby, Sweden (European patent no. 1383913, US patent no. 7,465,559) to determine the tentative cut-off value for each of the molecules tested. In the event of outlier values, a maximum of 1% (n=2 isolates) of extremely high diameter values were removed for analysis. For MICs, the ECOFFFinder tool (https://www.eucast.org/mic_distributions_and_ecoffs/, v2.1) was used to determine the tentative cut-off value for each molecule tested without removing any outliers. The temporary cut-off considered was the ECOFF set at 99% of the estimated wild-type population. Isolates resistant to at least three different antimicrobial families were considered multi-drug resistant (MDR).

Scattergrams were plotted to compare the concordance between the DD and MIC categorisation of isolates. The percentage of discrepancy (Pd) was calculated for each antimicrobial agent tested by both methods. For a value less than 5%, the classifications obtained with the two methods were considered as equivalent (**Table S2**).

Statistically significant differences in the distribution of wild-type (WT) and non-wild-type (NWT) isolates of different origins (clinical poultry (CP) versus non-clinical poultry (NCP) isolates) according to DD results were confirmed by the Chi^2^ test or Fisher exact test for five or fewer isolates, for a p-value less than 0.05

### Identification of gene mutations

Mutations in genes that may be involved in antimicrobial resistance were identified by multi-alignment of the corresponding proteins using MultAlin (12). Only mutations specific to NWT isolates were selected.

## RESULTS

### Determination of inhibition zone diameters

The inhibition zone diameters were determined for the 208 *E. cecorum* isolates using the DD method. For each DD assay, the results obtained for the reference strain and the inoculation density complied with EUCAST recommendations (data not shown).

The temporary cut-Off (CO_WT_) value was determined for 25 molecules (**Figure 2, Table S3**). The distribution of the diameters of four antimicrobials is shown in **Figure S1**, while WT and NWT isolates are presented in **Figure 2**. The heterogeneity of diameters for four β-lactams (ampicillin, amoxicillin, cefotaxime and ceftaroline) prevented the determination of CO_WT_ values.

**Figure 2:**
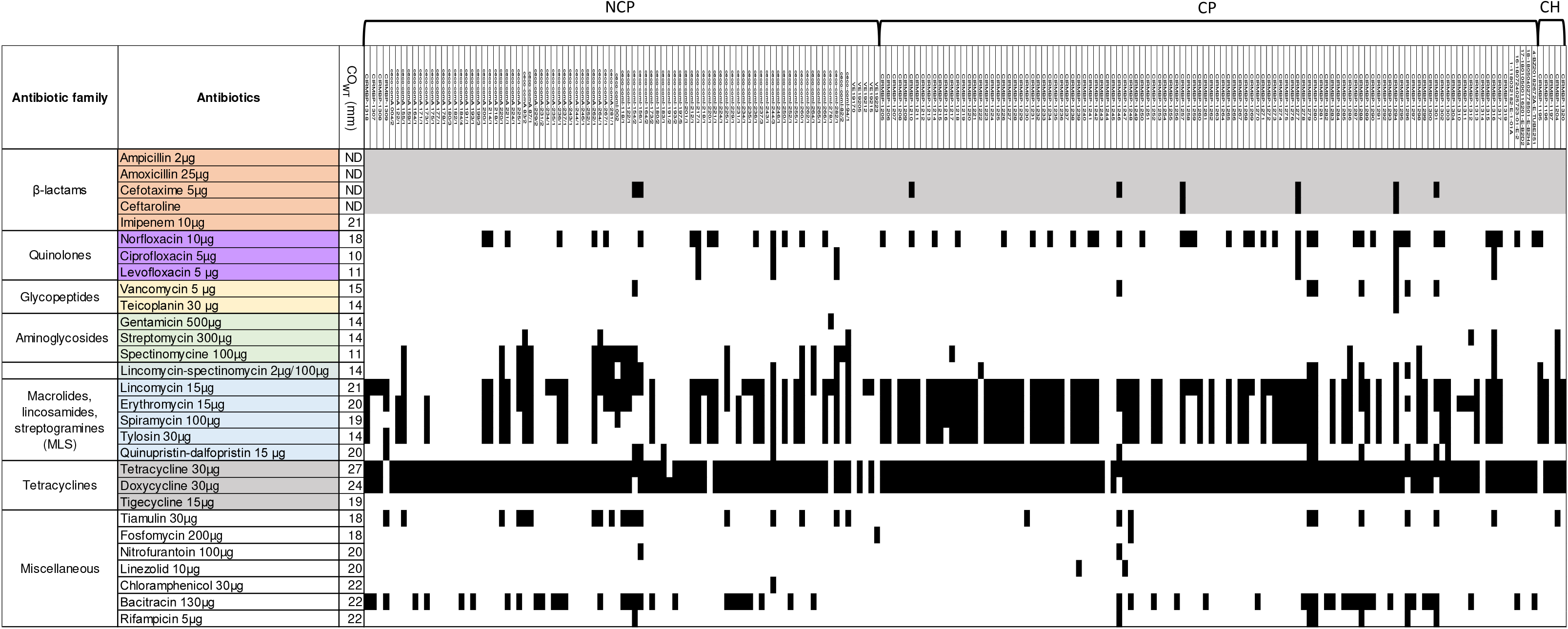
Heatmap of *E. cecorum* antimicrobial sensitivity determined by the disc diffusion method. Determination of WT (white), non-WT (black) and non-determined (grey) status for each antimicrobial. NCP: non-clinical poultry isolate, CP: clinical poultry isolate, CH: clinical human isolate. ND: CO_WT_ was not determined.

We tested several molecules for a few antimicrobial families: there was almost total agreement between DD results obtained for ciprofloxacin and levofloxacin, with respectively 97.6% and 97.1% of WT isolates, whereas 74.5% were categorised as WT for norfloxacin.

The concordance between spiramycin and tylosin was 98.6% and yielded similar rates of NWT isolates (48.1% to 47.6%). The classification according to DD for spiramycin and tylosin was similar to that obtained for erythromycin for respectively 93.3% and 92.8% of isolates. Most of the isolates tested using DD were resistant to tetracycline and doxycycline (respectively 95.7% and 93.8%, with an agreement of 98%) while none showed resistance to tigecycline.

Globally, there were no NWT isolates for imipenem and fewer than ten for ciprofloxacin (n=5) and levofloxacin (n=6), teicoplanin (n=1), vancomycin (n=8), gentamicin (n=1), streptomycin (n=5), fosfomycin (n=2), nitrofurantoin (n=2), linezolid (n=2), chloramphenicol (n=1) and rifampicin (n=7). Thirteen percent of the isolates were NWT for tiamulin. Of all the isolates, 43.3% were found to be MDR to up to eight different antimicrobial families, including glycopeptides. Only three isolates (1.4%) were susceptible to all the molecules tested.

Distributions of WT and NWT isolates according to their origin (**Table 1**) showed that identical rates of CP and NCP isolates were NWT for tetracycline and doxycycline, as were for bacitracin. There were significant differences in the ratios of WT and NWT isolates for erythromycin, as there were respectively 65% and 40% of NWT for CP and NCP (p= 0.00038); likewise for norfloxacin, where the corresponding figures for NWT were 32% and 18% (p= 0.025). On the other hand, for spectinomycin and lincomycin-spectinomycin, 21% of NCP isolates were NWT for both compared with 4% of CP isolates for spectinomycin (p=8.7×10^−5^) and 8% for lincomycin-spectinomycin (p= 6×10^−3^).

**Table 1:** Non-wild-type isolates according to their clinical origin.

|  | CP<sup>a</sup> (n=113) | NCP<sup>a</sup> (n=89) | P-value<sup>c</sup> |
| --- | --- | --- | --- |
| Antimicrobial | No. NWT<sup>b</sup> (%) | No. NWT<sup>b</sup> (%) |  |
| Norfloxacin | **36 (32%)** | **16 (18%)** | **0.025** |
| Ciprofloxacin | 2 (2%) | 3 (3%) | 0.66 |
| Levofloxacin | 3 (3%) | 3 (3%) | 1 |
| Vancomycin | 7 (6%) | 1 (1%) | 0.08 |
| Teicoplanin | 1 (1%) | 0 (0%) | 1 |
| Gentamicin | 0 (0%) | 1 (1%) | 0.44 |
| Streptomycin | 2 (2%) | 3 (3%) | 0.66 |
| Spectinomycin | **4 (4%)** | **19 (21%)** | **8.19x10<sup>-5</sup>** |
| Lincomycin/Spectinomycin | **9 (8%)** | **19 (21%)** | **6x10<sup>-3</sup>** |
| Lincomycin | 75 (66%) | 47 (53%) | 0.05 |
| Erythromycin | **74 (65%)** | **36 (40%)** | **3.8x10<sup>-4</sup>** |
| Spiramycin | **67 (59%)** | **29 (33%)** | **1.6x10<sup>-4</sup>** |
| Tylosin | **66 (58%)** | **29 (33%)** | **2.6x10<sup>-4</sup>** |
| Quinupristin/dalfopristin | 7 (6%) | 5 (6%) | 0.86 |
| Tetracycline | 111 (98%) | 83 (93%) | 0.14 |
| Doxycycline | 108 (96%) | 82 (92%) | 0.3 |
| Tigecycline | 0 (0%) | 0 (0%) | 1 |
| Tiamulin | **9 (8%)** | **18 (20%)** | **0.01** |
| Fosfomycin | 1 (1%) | 1 (1%) | 1 |
| Nitrofurantoin | 1 (1%) | 1 (1%) | 1 |
| Linezolid | 2 (2%) | 0 (0%) | 1 |
| Chloramphenicol | 0 (0%) | 1 (1%) | 1 |
| Bacitracin | 26 (23%) | 29 (33%) | 0.12 |
| Rifampicin | 6 (5%) | 1 (1%) | 0.14 |

<sup>a</sup> CP for clinical poultry and NCP for non-clinical poultry isolates
<sup>b</sup> WT for wild-type isolates and NWT for non-wild-type isolates according to the disc diffusion method
<sup>c</sup> P-value calculated with Chi<sup>2</sup> or Fisher exact test if n≤5. Significant p-values are in bold.

### Determination of minimal inhibitory concentrations (MICs)

For each MIC determination assay, the results obtained for the reference strain complied with EUCAST recommendations (data not shown) (EUCAST, 2019). The distributions of the MICs of 23 antimicrobials and their corresponding MIC_50_ and MIC_90_ values are displayed in **Figure 1**. MIC values above the tested range were observed for seven antimicrobials (vancomycin, teicoplanin, tylosin, tetracycline, doxycycline, bacitracin and avilamycin). For glycopeptides, doxycycline and bacitracin, only one, two or three strains were concerned. The greatest proportion of isolates with MIC values above the tested range were found for tetracycline (n=128, 61.5%), tylosin (n=54, 26%) and avilamycin (n=41, 19.7%). For teicoplanin, tetracycline and rifampicin, the MIC_50_ and MIC_90_ values were equal (respectively 0.125 mg/L, 256 mg/L and 0.125 mg/L). For ten (ampicillin, amoxicillin, vancomycin, gentamicin, doxycycline, tigecycline, chloramphenicol, linezolid, avilamycin and daptomycin) of the tested antimicrobials, the values between MIC_90_ and MIC_50_ differed by only one dilution step. CO_WT_ values were calculated for 17 antimicrobial agents for the complete *E. cecorum* dataset (**Figure 2**). No value could be computed for beta-lactams nor for spectinomycin due to the heterogeneity of the population distribution. For tetracycline, doxycycline and avilamycin, CO_WT_ values were determined through a graphical representation due to the number of NWT strains. For seven antimicrobial agents, the percentages of NWT isolates were high to extremely high (tylosin (38%), spiramycin (38.5%), lincomycin (42.8%), erythromycin (44.2%), tetracycline (77.9%), doxycycline (81.7%) and avilamycin (90.9), whatever their origin.

### Comparison between MICs and inhibition zone diameters for *E. cecorum*

**Table S2** highlights the difference in interpretation between DD and BMD methods. For nine antimicrobial agents, the Pd values were lower than 5% (levofloxacin 4.8%, vancomycin 3.85%, teicoplanin 0.5%, gentamicin 1%, quinupristin-dalfopristin (QDF) 2.9%, tigecycline 0.0%, chloramphenicol 0.5%, linezolid 1% and rifampicin 2.4%), meaning that there were very few differences in WT/NWT categorisation between the methods. In contrast, nine antimicrobial agents yielded Pd values higher than 5% (ciprofloxacin 7.7%, streptomycin 9.6%, lincomycin 24.5%, erythromycin 16.4%, spiramycin 14.4%, tylosin 15.4%, doxycycline 14.9%, tetracycline 17.8%, and bacitracin 17.8%). For macrolides, lincosamides and streptogramins (MLS), tetracyclines and bacitracin, CO_WT_ values determined by the DD method gave more NWT strains than BMD did. Conversely, for ciprofloxacin and streptomycin, CO_WT_ values determined by the BMD method gave more NWT strains than DD.

### Consistent gene content and expression of resistance phenotypes

In our previous genomic study isolates (J. Laurentie, V. Loux, C. Hennequet-Antier, E. Chambellon, J. Deschamps, A. Trotereau, S. Furlan, C. Darrigo, F. Kempf, J. Lao, M. Milhes, C. Roques, B. Quinquis, C. Vandecasteele, R. Boyer, O. Bouchez, F. Repoila, J. Le Guennec, H. Chiapello, R. Briandet, E. Helloin, C. Schouler, I. Kempf, and P. Serror, submitted for publication), antimicrobial resistance genes were identified on the genomes of 118 isolates and correspondences between their genomic content and their resistance phenotype are presented below (**Table S4**).

The 114 tetracycline-NWT isolates with sequenced genomes carried at least one tetracycline resistance gene. The *tet*(M) *tet*(L) association was the most frequent (n=61), followed by *tet*(M) gene only (n=47), the associated *tet*(M) *tet*(O) genes (n=2), *tet*(M), *tet*(L) and *tet*(O) (n=1), *tet*(M) and *tet*(44) (n=1), the *tet*(O) gene (n=1) and the *tet*(L) gene (n=1). Thus, the ribosomal protection gene *tet*(M) is the most frequent tetracycline resistance gene (n=112) in our *E. cecorum* isolates, as reported for *E. cecorum* (9) and other enterococci [16]. No tetracycline-resistance gene was detected in the genomes available for the four tetracycline-WT isolates. As expected, most of our *erm*-positive isolates were NWT for erythromycin (78/80, 97.5%) and lincomycin (77/80, 96.3%). Four isolates bearing *mefA* and *lnuC* were NWT for lincomycin but not the other tested MLSs. The four isolates possessing the *ermG, mefA, msrD* and *lnuC* genes were NWT for erythromycin, but only one was also NWT for lincomycin, spiramycin and tylosin. Six of the eight QDF-resistant isolates for which genomic sequences were available carried the *ermB* or *ermG* gene, but no other acquired resistance mechanisms could be detected in the remaining two. Two isolates with an *ermB* gene were found WT for macrolides. These diverse discrepancies between MLS phenotypes and genotypes were not investigated further, but could result from the poor or lack of expression of the encoded proteins, and/or the presence of unknown resistance genes or mutations. Acquired resistance genes were identified to explain resistance to gentamicin, streptomycin or spectinomycin in six aminoglycoside-resistant isolates. The *spc* (or *ant*(9)-Ia) gene was detected in two spectinomycin-resistant isolates. Four isolates contained *aadE* (*ant*(6)-Ia), an aminoglycoside-nucleotidytransferase conferring resistance to streptomycin, and were found resistant to this antimicrobial according to both BMD and DD methods. Two isolates from the United States have been reported to carry the *aph*(3’)-III gene conferring resistance to kanamycin (9), but this aminoglycoside was not tested. According to the DD or BMD method, 26.4% or 16.3% of isolates were NWT for bacitracin. Among the sequenced isolates, 27 were found to be NWT using DD and 24 of these NWTs had a bacitracin resistance operon, with *uppP2, bcrB, bcrA* and *bcrR* genes. This operon was also detected in only four of the 91 bacitracin-WT sequenced isolates. With regard to glycopeptides, seven isolates were classified as NWT for vancomycin according to DD. One of them was also NWT for teicoplanin and a *vanA* operon was detected in its genome. No other glycopeptide resistance genes were detected.

For fluoroquinolones, we looked in the 118 available genome sequences for chromosomal mutations that could explain resistance as they may target DNA gyrase and topoisomerase IV. The three ciprofloxacin- and levofloxacin-NWT sequenced isolates harboured a mutation in the GyrA quinolone resistance-determining region (either S83Y or S83V), only detected in these genomes. Moreover, a mutation in ParC (S82I) was also highlighted in 24/35 of norfloxacin-NWT isolates.

## DISCUSSION

For the first time, the CO_WT_ values for 25 out of 29 and 20 out of 23 molecules tested using the DD or BMD methods respectively were calculated with the EUCAST method adapted to *E. cecorum*, (or in three cases by visual inspection for BMD) the main exception being for β-lactams, which have a heterogeneous distribution. The availability of tentative ECOFF values is the first step in monitoring the AMR of a bacterial species. Our data are based on results from one laboratory only; however, they concern a variety of commensal and clinical isolates collected over more than 37 years. They will no doubt contribute to the definition of official interpretative criteria for *E. cecorum*, a major poultry pathogen. Data from other countries relative to multiple and diverse sources and a large number of isolates are now needed to set recognised ECOFFs with the expected precise estimates (13).

This work reveals that the concentration criteria currently used to classify *E. cecorum* isolates, based on the breakpoints recommended for enterococci or ECOFF values for *E. faecalis* and *E. faecium*, do not always properly distinguish WT from non-WT, while the ECOFF values for *S. pneumoniae* coincide more frequently (**Table 2**). Part of this difference is probably related to the similar growth conditions under CO_2_ 5% for *E. cecorum* and *S. pneumoniae*. Overall, discrepancies between the distribution of MICs and CO_WT_ or ECOFF values for *E. cecorum* and *S. pneumoniae, E. faecium* or *E. faecalis* were observed for aminoglycosides, tigecycline, avilamycin and bacitracin. Indeed, *E. cecorum* seems susceptible to moderate concentrations of aminoglycosides, and the CO_WT_ determined for streptomycin (32 mg/L) and gentamicin (4 mg/L) are lower than the ECOFF for *E. faecium* (respectively 128 mg/L and 32 mg/L) or *E. faecalis* (512 and 64 mg/L). Currently there are no ECOFF values for these antimicrobials for *S. pneumoniae*. The calculated CO_WT_ for tigecycline is 4 mg/L, which is much higher than the ECOFF for *S. pneumoniae* (0.125 mg/L) or *E. faecium* (0.25 mg/L).

**Table 2.** Comparison of *E. cecorum* CO_WT_ values with *S. pneumoniae, E. faecalis, E. faecium* ECOFF or criteria previously used for *E. cecorum*.

| Antimicrobial | Broth micro-dilution<br>MIC (mg/L) |  |  |  |  | Disc diffusion<br>Inhibition zone diameter (mm) |  |  |  |
| --- | --- | --- | --- | --- | --- | --- | --- | --- | --- |
|  | CO<sub>WT</sub><br>*E.*<br>*cecorum* | ECOFF<br>*S. pn*<sup>a</sup> | ECOFF<br>*E.*<br>*faecalis* | ECOFF<br>*E.*<br>*faecium* | Other<br>criteria<sup>b</sup> | CO<sub>WT</sub><br>*E.*<br>*cecorum* | ECOFF<br>*S.pn* | ECOFF<br>*E.*<br>*faecium* | ECOFF<br>*E.*<br>*faecalis* |
| Ampicillin | ND | 0.06 | 4 | 8 | 0.25<sup>d</sup><br>(AMX) | ND | (28) | 10 | 10 |
| Avilamycin | 1 |  | 8 | 16 |  |  |  |  |  |
| Bacitracin | 4 |  | (32)<sup>f</sup> | 32 |  | 22 |  |  |  |
| Chloramphenicol | 8 | 8 | 32 | 32 | 16<sup>e</sup> | 22 | 21 |  |  |
| Ciprofloxacin | 4 | 4 | 4 | 8 | 0.5<sup>e</sup><br>(ENR<sup>g</sup>)<br>2<sup>e</sup> | 12 | (18) |  |  |
| Daptomycin | 0.25 | (0.5)<sup>f</sup> | 4 | 8 | 4<sup>e</sup> |  |  |  |  |
| Doxycycline | 1 | 0.5 | 0.5 | 0.5 |  | 24 |  |  |  |
| Erythromycin | 0.5 | 0.25 | 4 | 4 | 0.5<sup>d</sup><br>4<sup>e</sup> | 21 | 22 |  |  |
| Fosfomycin |  |  | 128 |  |  | 18 |  |  |  |
| Gentamicin | 4 |  | 64 | 32 | 4<sup>c</sup><br>250<sup>d</sup> | 15 |  | 8 | 8 |
| Imipenem |  | 0.016 | 4 | 4 |  | 21 |  | 21 | 21 |
| Levofloxacin | 2 | 2 | 4 | 4 |  | 13 | 19 |  |  |
| Linezolid | 8 | 4 | ID | 4 | 4<sup>e</sup> | 21 | (22) | 19 | 19 |
| Nitrofurantoin |  |  | (32)<sup>f</sup> | 256 |  | 21 |  |  | 15 |
| Norfloxacin |  |  |  |  |  | 18 | 12 |  | 12 |
| Quinupristin-dalfopristin | 4 |  |  |  | 2<sup>e</sup> | 20 |  |  |  |
| Rifampicin | 1 | (0.125)<sup>f</sup> |  |  |  | 22 | 25 |  |  |
| Streptomycin | 32 |  | 512 | 128 | 512<sup>c</sup><br>500<sup>e</sup> | 15 |  | - | - |
| Teicoplanin | 1 | 0.25 | 2 | 2 |  | 14 | (18)<sup>f</sup> | (16)<sup>f</sup> | (16)<sup>f</sup> |
| Tetracycline | 2 | 1 | 4 | 4 | 4<sup>c</sup><br>8<sup>e</sup> | 27 | 25 |  |  |
| Tigecycline | 4 | 0.125 | 0.25 | 0.25 | 0.25<sup>e</sup> | 19 |  |  | 18 |
| Tylosin | 8 |  |  |  | 2<sup>d</sup><br>16<sup>e</sup> | 15 |  |  |  |
| Vancomycin | 2 | 1 | 4 | 4 | 16<sup>e</sup> | 15 | (16)<sup>f</sup> | 12 | 12 |

<sup>a</sup> *S. pn* : *Streptococcus pneumoniae*
<sup>b</sup> Criteria used by Borst et al. (Borst et al., 2012) based on <sup>c</sup> breakpoints for enterococci (Clinical Laboratory Standards Institute, 2010) or <sup>d</sup> stratification using the overall median concentration into susceptible (below the median) and non-susceptible (above the median) or <sup>e</sup> criteria used by Jackson et al. (Jackson et al., 2015).
<sup>f</sup> EUCAST tentative ECOFF
<sup>g</sup> ENR for enrofloxacin
ECOFF values are obtained from the EUCAST website (mic.eucast.org/search/)

Both the MIC_50_ and MIC_90_ for tigecycline are low (respectively 0.5 and 1 mg/L) and all 208 isolates were classified as WT according to both DD and BMD methods. Our MIC results for avilamycin yielded a MIC_50_ of 64 mg/L and a CO_WT_ value of 1 mg/mL compared to ECOFF values of 8 mg/L or 16 mg/L respectively for *E. faecalis* or *E. faecium*, resulting in a percentage of avilamycin-NWT *E. cecorum* of 91.0%. The calculated CO_WT_ for bacitracin was also three double dilutions lower than the ECOFF value for *E. faecium*. The size of the NWT population may also bias the determination of the CO_WT_. We could not calculate the CO_WT_ of beta-lactams for *E. cecorum*. The MIC_90_ values for ampicillin and amoxicillin were lower (0.25 mg/L and 0.125 mg/L respectively) than the MICs reported for *E. faecalis* or *E. faecium* (https://mic.eucast.org/), a feature of *E. cecorum* already reported for penicillin by Jung et al. (4). Using an amoxicillin breakpoint of 0.25 mg/L like Borst et al. (6) we managed to classify nine out of 208 isolates as amoxicillin-resistant (4.0%) in contrast to the 26.3% NWT rate reported by the authors. This observation suggests a low rate of amoxicillin resistance in our collection.

A comparison of the interpretations between the DD and BMD methods used and the gene content of the sequenced isolates shows that the DD method is more consistent with the gene content. For erythromycin, for example, using the DD method, 82/84 NWTs have an *erm-* gene and only two WT isolates carry one. Conversely, according to the BMD method, 13 isolates of the 47 considered as WT carry an *erm* gene and 66/71 of the NWTs carry one. The same is true, to a lesser extent, for tetracycline: 114/114 NWTs classified by the DD method carried tetracycline resistance genes while 110/114 are considered NWTs by the BMD method and carry at least one *tet* gene. These different observations suggest that under the conditions tested, the DD method and the corresponding calculated CO_WT_ value are more accurate in determining the resistance of *E. cecorum*.

Despite a significant decrease in the use of antimicrobials in animals during the last ten years in France, Europe and the USA, tetracyclines and penicillins are still used extensively (14). The prevalence of NWT isolates for tetracycline was extremely high, as previously observed for enterococci from French broilers (15). High rates have also been reported for *E. cecorum* in other countries (70%) (4), as well as for *E. faecium* (80.3%) and *E. faecalis* (78.5%) (16). The prevalence of NWT isolates for erythromycin, a 14-membered macrolide, was quite high (44.7% for BMD and 54.8% for DD), although lower than in data previously reported for *E. cecorum* (70% (4)), and similar to those reported for European broiler isolates of *E. faecium* (57.0% %) and *E. faecalis* (56.6%) (16). CP isolates were more frequently resistant to MLSs (erythromycin, spiramycin and tylosin) than NCP ones. This difference has already been observed in other studies (6, 7, 9, 17). Indeed, macrolides are used for therapy for poultry, which may explain the prevalence of resistance in clinical isolates. A slightly higher number of *E. cecorum* strains were found to be NWT for lincomycin (60.6% according to DD), but with a MIC_50_ of 1 mg/L, this species does not seem naturally resistant to lincosamides, unlike several other *Enterococcus* species (e.g. *E. faecalis, E. avium, E. gallinarum, E. casseliflavus*). The association of lincomycin (66% of NWTs in CP isolates) and spectinomycin (4% of NWTs in CP isolates) highlights the benefits of lincomycin, as only 8% of CP isolates are NWT for lincomycin-spectinomycin. This susceptibility may explain the reported use in Germany of the antimicrobial association during the first week of life to control *E. cecorum* isolates (18).Similarly, the MIC_90_ of quinopristin-dalfopristin was low (2 mg/L) for our *E*. cecorum isolates, and precludes an intrinsic resistance to these compounds, unlike *E. faecalis*. Indeed, only 6.2% of isolates were classified as NWT for quinopristin-dalfopristin. It should be remembered that high occurrences of quinopristin-dalfopristin-NWT *E. faecium* were observed in poultry in France and other European countries several years ago, probably due to the use of virginiamycin as a growth promoter in Europe up to 1999 (15, 19). Unfortunately, this quinopristin-dalfopristin resistance apparently persisted in *E. faecium* up to recent observations, maybe as a result of co-selection by other antimicrobials used for therapy (19). Different resistance genes to MLS (*ermB, ermG, lnuB, merfA, msrD, linB, lnuB, lnuC* and *lnuD* have already been identified by Sharma et al. in *E. cecorum* genomes and associated with resistance phenotypes (9).

Avilamycin is an oligosaccharide antimicrobial used as a growth promoter among poultry in the European Union up to 2006. A link between the consumption of avilamycin and resistance of *E. faecium* to this antimicrobial has been demonstrated in the literature (20, 21). In France, the percentages of broilers or turkeys with avilamycin-resistant *E. faecium* were respectively 8.3% and 21.8% in 2007 (22). Thus the high level of resistance of *E. cecorum* (90.8% of NWT, and a MIC_50_ of 64 mg/L) to an antimicrobial that is no longer used is rather unexpected, but we could not detect the previously described avilamycin resistance mechanisms, i.e. the *emtA* gene or mutations in L16 (20).

Very few isolates were classified as NWT for chloramphenicol, fosfomycin and nitrofurantoin. A low prevalence of chloramphenicol resistance has also been reported in other studies for *E. cecorum* or *E. faecium* (4, 23). Since 1994, chloramphenicol is forbidden in Europe for food-producing animals, but other phenicols (e.g. florfenicol) are authorised, mainly for pigs and cattle, explaining more frequent chloramphenicol-NWT *E. faecalis* in these productions (23). Fosfomycin has rarely been tested against poultry enterococci. According to Schwaiger et al. (24) about 5% of *E. faecalis* from cloacal swabs of organically and conventionally kept laying hens were resistant to fosfomycin. In enterococci, fosfomycin can be related to mutations in the target enzyme MurA, or enzymatic modification of fosfomycin, linked to the acquisition of transferable *fosB* genes or the high expression of the *fosX* gene (25, 26). Consistently with Suyemoto et al. (27) who detected no nitrofurantoin resistance among 32 *E. cecorum* strains, the prevalence of this resistance in our collection is below 1%.

Prevalence of *E. cecorum* resistance to critically important antimicrobials in human medicine is quite low, confirming our prediction from genome analysis isolates (J. Laurentie, V. Loux, C. Hennequet-Antier, E. Chambellon, J. Deschamps, A. Trotereau, S. Furlan, C. Darrigo, F. Kempf, J. Lao, M. Milhes, C. Roques, B. Quinquis, C. Vandecasteele, R. Boyer, O. Bouchez, F. Repoila, J. Le Guennec, H. Chiapello, R. Briandet, E. Helloin, C. Schouler, I. Kempf, and P. Serror, submitted for publication). Low rates of linezolid NWT isolates agree with the low detection levels among broilers in *E. faecium* and *E. faecalis* in 2012-2013 (15, 23). However, recent studies on animal samples inoculated onto linezolid-supplemented media revealed that linezolid-resistant enterococci (or staphylococci) may be frequently present although not detected when analysing only strains from the dominant enterococci population isolated on non-supplemented media (28). Linezolid has never been used in animal production, but antimicrobial resistance mechanisms evidenced in animal isolates (e.g. *cfr, optrA* and *poxtA* genes) could explain the co-selection of the described linezolid-resistant strains by the use of other commonly used antimicrobials (e.g. tetracyclines, florfenicol in pigs) (29). Daptomycin is not used in animal production either, but is an option for treating infections with multidrug resistant or vancomycin-resistant enterococci (VRE) in human patients. In line with our very low rate of daptomycin-NWT isolates, most isolates of *E. faecium* and *E. faecalis* and all *E. hirae, E. durans* and *E. casseliflavus* isolates from healthy cattle, pigs and chickens in nine EU countries yielded susceptible isolates (19). Likewise, no daptomycin-resistant *E. cecorum* strain was present in the collection analysed by Suyemoto et al. in the USA (27). Tigecycline is a glycylcycline developed to overcome tetracycline resistance mechanisms. The activity of this last resort antimicrobial for human medicine appears to have been preserved up to now in enterococci from animals, as the absence of tigecycline resistance in our *E. cecorum* isolates is in line with the results described for *E. faecium, E. hirae, E. durans* and *E. casseliflavus* isolates of animal origin (4). The low gentamicin resistance rates are also in line with the findings reported for *E. faecium* and *E. faecalis* from animals in Europe (4) but higher levels were sometimes observed for *E. cecorum* (6, 17). The prevalence of vancomycin resistance was low, as observed in recent studies of *E. cecorum* (4). Noteworthy, the *vanA* operon has already been detected in an *E. cecorum* isolate in Japan (30). Strikingly, although rifampicin is not used in poultry production in France, seven isolates were classified as NWT. Resistance to rifampicin in enterococci is usually associated with mutations in the RNA polymerase gene (*rpoB*), but no mutations specific to *E. cecorum* NWT isolates were detected.

## Conclusion

For the first time, the tentative cut-offs for 25 and 20 antimicrobials were determined using respectively DD and BMD methods on a large collection of *E. cecorum* with clinical and non-clinical origins. Further experiments could confirm or refine these cut-off values. The DD method seems more appropriate to determine resistance thresholds for this bacterium, as it is more in line with the genetic content. Resistance to medically important antimicrobials (imipenem, vancomycin, gentamicin, tigecycline or linezolid) is rare. Mutated GyrA (S83Y or S83V) and ParC (S82I) may explain resistance to some fluoroquinolones. Moreover, this study revealed that while some resistances (tetracycline, erythromycin, etc.) are still prevalent, strains of clinical origin showed significant levels of sensitivity for antimicrobials authorised for poultry.

## ACKNOWLEDGEMENTS

The authors are grateful to Vincent Cattoir (CHU Rennes, France) for advice in antimicrobial selection, to Avipole (Ploufragan, France), J. Le Guennec (Labofarm, Loudéac, France) and Labocea35 (Fougères, France) who provided caecal contents to isolate commensal isolates, and to Bioscand AB (Täby, Sweden) for providing the NRI tool.

## Supplementary material

**Table S1:** Isolates used in the study.

**Table S2:** Scattergram comparing MICs and inhibition zone diameters.

**Table S3:** CO_WT_ values, inhibition zone diameters and interpretations for the disc diffusion method.

**Table S4:** Gene content, mutations and phenotype of resistance in 118 *E. cecorum* isolates.

**Figure S1**: Distribution of inhibition zone diameters and NRI method for CO_WT_ determination. A. Quinupristin-Dalfopristin; B. Erythromycin; C. Vancomycin; D. Amoxicillin. The orange dotted line represents the CO_WT_ value determined. No CO_WT_ could be determined for amoxicillin.

